# Vertical transmission of Zika virus in orally-infected *Aedes aegypti* produces infectious adult progeny

**DOI:** 10.1101/375964

**Authors:** Genevieve Comeau, Robert A. Zinna, Taylor Scott, Kacey Ernst, Kathleen Walker, Yves Carrière, Michael A. Riehle

## Abstract

Vertical transmission, or pathogen transfer from mother to offspring, can facilitate persistence of emerging arboviruses, such as Zika virus (ZIKV), in mosquito populations. Understanding vertical transmission and the different environmental and temporal conditions that affect it is important to assess whether new outbreaks could occur without reintroduction of the virus. To determine the rate of vertical transmission for ZIKV, *Aedes aegypti* females were fed on ZIKV infected blood, maintained under three temperature conditions (27°C, 30°C, and 33°C), and allowed to oviposit three times. Progeny were tested for virus presence at 3, 7, and 14 days after adult emergence. The overall vertical transmission rate was 6.5% (3.9 - 9.9). Vertical transmission was observed across all maternal temperature conditions and was detected in adult progeny as young as 3 days and as late as 14 days post-emergence. In total, 3.4% (1.6 - 6.2) of adult progeny produced saliva with detectable ZIKV, indicating their capacity to transmit ZIKV to humans. To our knowledge, this is the first evidence that vertical transmission occurs from orally-infected female *Aedes aegypti* to their adult progeny at a range of temperatures, and proof that Zika virus can persist in the saliva of those progeny throughout their lifetimes. These results suggest that the virus may be maintained in *Ae. aegypti* populations without a vertebrate host, allowing for human infections to occur without consistent re-introductions of ZIKV.

**Author Summary:** In 2015, Zika virus spread to over 50 countries. However, it is not known whether the virus persisted in the outbreak areas or became locally extinct. One way mosquito-borne viruses, like Zika, could become established is by transferring directly between mosquito generations rather than circulating between mosquitoes and humans. This is known as vertical transmission, and happens when the virus infects the developing eggs of infected maternal mosquitoes. As with other mosquito-borne diseases, like dengue, in order to infect humans the virus must be present in the saliva of infected mosquito progeny during blood feeding. We found vertical transmission occurred throughout the infected mother’s reproductive lifetime and across a range of temperature conditions. Vertically infected progeny had Zika virus in their saliva as early as three days after adult emergence, implying that they could infect a person even during their first bloodmeal. Importantly, this work indicates that Zika virus could establish itself in the mosquito population even when human to mosquito transmission is not actively occurring.

## Introduction

Zika virus (ZIKV) is a mosquito-borne flavivirus that emerged as a significant human pathogen in French Polynesia and has since spread to over 50 countries. (*1*) While asymptomatic in approximately 80% of cases, ZIKV infection can cause adverse pregnancy outcomes, such as Zika congenital syndrome (2), and serious neural complications in children and adults including Guillain-Barre and fatal encephalitis. *(3)* It is primarily transmitted to humans through the bite of the yellow fever mosquito *Aedes aegypti*, although other mosquito species are also competent. (*4*) Following the pandemic in 2015, 2.2 billion people live in areas that reported active ZIKV transmission. (*5*) A better understanding of how ZIKV becomes permanently established after introduction to these new locales is needed to assess risk and prevent disease.

Arthropod-borne viruses, or arboviruses, can become established in a geographic region through several mechanisms, including 1) regular viral reintroduction and circulation across geographies, 2) established human or non-human reservoirs, and 3) virus maintenance in the vector population. (6) Maintenance in the vector population keeps viruses prevalent when there are few human or other hosts, such as between outbreaks (7). It also helps arboviruses, such as West Nile, Ross River, and Sindbis, survive cold temperatures and low vector population numbers during harsh winter months. *(8, 9)* Vertical transmission, or direct pathogen transfer from mother to offspring, is one path to arboviral maintenance in vector populations. Vertical transmission of other *Ae. aegypti-borne* viruses such as dengue, chikungunya, yellow fever, and West Nile occurs in both the laboratory and field for *Ae. aegypti. (10-13)* Interestingly, vertical transmission occurs at a higher rate in *Aedes* than other genera of mosquitos. (*7*) If Zika is also vertically transmitted, it may increase the risk of becoming established in a given area and causing future outbreaks without reintroduction.

Vertical transmission of ZIKV alone is not sufficient for another outbreak to occur. To infect a human, virus must be present in the saliva of the vertically-infected progeny. Such progeny would then bypass the extrinsic incubation period (EIP), which is the time between the ingestion of a viremic blood meal by a female mosquito and the time when virus is present in the saliva. Estimates of the EIP of ZIKV range from 4-10 days *(14, 15)* and this EIP could potentially limit ZIKV spread due to the limited lifespan of *Ae. aegypti* in the field *(16)*. Furthermore, vertically infected mosquitoes do not need to obtain an initial ZIKV infected blood meal, leading to increased infection rates and younger infected mosquitoes. If the saliva of vertically-infected mosquitoes is infected with ZIKV earlier than horizontally-infected mosquitoes, they would have more opportunities to transmit the virus. However, the capacity of vertically-infected mosquitos to develop infectious saliva has not been evaluated.

Previous studies have implicated ZIKV as vertically transmitted, but evidence is limited by study conditions and endpoints. One study demonstrated a low rate, (1/290) adults, of vertical transmission in adult progeny. However, virus was introduced intrathoracically into the maternal mosquitoes, which bypasses midgut invasion and escape barriers and could impact the rate of infection. *(17)* A subsequent study demonstrated higher rates of vertical transmission in immature mosquitos, 1/84 larvae, but this rate might not accurately reflect infection status following adult eclosion and does not provide insights into whether ZIKV infected adults have virus in their saliva. *(18)* Neither study examined how vertical transmission varies under different environmental conditions, though factors such as temperature and humidity conditions can strongly influence arboviral transmission dynamics. *(19-22)* Gonotrophic cycle, or number of egg batches laid, and adult mosquito age also affect vertical transmission rates of arboviruses. (*7, 23*) To date, no studies have investigated the rate of ZIKV vertical transmission at multiple temperature conditions, gonotrophic cycles, or whether infection persists during the lifetime of adult progeny.

The objective of this study, therefore, was to quantify the rate of ZIKV vertical transmission to adult offspring of orally infected female *Ae. aegypti* mothers across multiple maternal temperatures, gonotrophic cycles, and adult progeny ages. As many countries reported active ZIKV transmission after the 2015 outbreak, the results of this study will help inform public health policy, surveillance strategies, and potential interventions.

## Methods

### Mosquito Rearing

Mosquitoes originated from an established lab colony of UGAL strain *Ae. aegypti*. and were reared in an ACL2 insectary at 27°C, 75% RH, and 16:8 light:dark photoperiod cycle. Larval mosquitoes were reared at a density of 100-150 larvae per liter of water and fed ground cat chow. Pupae were transferred to emergence cages with a density of 50 mosquitoes/cage. Adult mosquitoes were provided 10% sucrose *ad libitium* via soaked cotton balls. Sucrose soaked cotton balls were replaced with water soaked cotton balls 24 hours before bloodmeals to encourage feeding.

### Cell Culture and Virus Propagation

Vero cells were cultured in DMEM supplemented with 10% FBS and incubated at 37°C and 5% CO2. At 80% confluence, cells were either split 1:5 or infected with ZIKV. During infection old culture media was removed, then 2 mL of fresh media was mixed with virus (ZIKV strain PRVABC59, ATCC) and diluted to a multiplicity of infection of 10. After dilution, cells were incubated for 1 hour at 37°C and 5% CO2, rocking the flask every 15 min. After 1 hour, the infectious media was removed and fresh media added to the flask. After new media was added, cells were incubated for 96 hours at 37°C and 5% CO2 until 90% cytopathic effect was attained. Cell culture media was then pipetted into a 15 mL tube and centrifuged at 300 x g for 10 minutes *(24)*. Viral supernatant was transferred to a new 15 mL tube, mixed to 20% volume/volume with fetal calf serum, and stored at −80°C until use.

### Plaque Assay

Plaque assays were conducted based on protocols provided by VIRAPUR and those developed by *Agbulos et al* (*25*) to determine the ZIKV stock viral titer prior to oral infections, with slight modifications. ZIKV stock titers were also assessed using qPCR (see ZIKV qPCR methods below). For plaque assays, Vero cells were plated in 6-well plates and incubated overnight. Media was removed from the cells and viral stock diluted in serum-free DMEM from 10^-2^ to 10^-6^. The ZIKV dilutions were allowed to adsorb onto cells for 1 hour, rocking every 15 minutes to distribute virus among the cells. Following adsorption, infectious media was removed and 3 mL of DMEM mixed with 4% agarose was overlaid onto the cells. Overlaid Vero cells were incubated for 5 days at 37°C and 5% CO2, then stained with 0.1 volume of 5mg/mL MTT and incubated for a minimum of two hours before final imaging and quantification.

### ZIKV qPCR standard generation and viral qPCR quantification assay

A region of the envelope protein of ZIKV strain PRVABC59 (833 bp) was generated to quantify viral load using specific primers (Forward: ATCTAGAAGAGCCGTGACGC; Reverse: CTGAAAAGTCAAGGCCTGTC) which were designed to flank the qPCR amplicon primers developed by Franz et al (26): Viral RNA was extracted from infected media using Trizol as follows. 100 uL of viral supernatant was added to 500 uL of Trizol. Chloroform (50 ul) was added and the sample incubated for 10 minutes at room temperature, followed by centrifugation at 4°C and 14,000g for 15 minutes. The aqueous phase (150 ul) was removed and incubated for one min at room temperature with 50 ul of isopropanol, then centrifuged at 4°C and a 14000g for 15 minutes. Ethanol (400 ul, 80%) was added to each sample and incubated at room temperature for 20 min to evaporate, then resuspended in DEPC water. The isolated ZIKV total RNA was converted into cDNA using Applied Biosystem’s High-Capacity cDNA kit (ThermoFisher, Waltham, MA). The region of interest was subsequently amplified using PCR (GoTaq, Promega, Madison, WI) and the product purified using the QIAquick Gel extraction kit (Qiagen, Hilden, Germany). This product was ligated into the pGEM-T Easy Vector (Promega, Madison, WI), and then transfected into competent *E. coli (JM109)*. After transfected colonies were selected, the plasmid was purified using the QIAprep Spin Miniprep kit (Qiagen, Hilden, Germany) and sequenced to verify the ZIKV amplicon. By modifying the forward primer above to include a T7 flanking sequence, the validated plasmid was used as a template for PCR to generate cDNA. The new PCR product was purified and used as a template to generate pure ssRNA with the MEGAscript T7 RNA synthesis kit (Ambion, ThermoFisher, Waltham, MA) according to manufacturer protocols. The ssRNA standard was quantified using a Nanodrop 2000 (ThermoFisher, Waltham MA), and the concentration of RNA was used to calculate the number of ssRNA copies (N) in the standard. The quantified ssRNA was diluted to generate a standard curve for absolute qPCR quantification with a concentration range of 2.13E+8 copies – 2.13E+4 copies that was incorporated into each plate.

The following primers and Taq-man probes for the detection of ZIKV with qPCR methodology were used; Forward: CCGCTGCCCAACACAAG, Reverse: CCACTAACGTTCTTTTGCAGACAT, Probe: FAM-CTYAGACCAGCTGAAR-BBQ (16). ZIKV qPCR quantification was performed using the iTaq Universal Probes One-Step Kit (Bio-Rad, 172-5140) according to manufacturer protocol with a 10 ul reaction volume. RNA extracted from uninfected female *A. aegypti* was included in each plate as a negative control. Reactions were run on an Eppendorf RealPlex2 Mastercycler for 10 min at 50°C, 3 min at 95°C., 95°C for 15, then 60°C for 30 sec. Samples were considered positive for ZIKV if amplification was detected at or before a C_t_, cycle threshold, value of 35. This number is highly conservative, as the CDC uses a C_t_ cut off value of 38. *(27)*

### Preparation of Infectious Bloodmeal and Oral infection of Mosquitoes

PRVABC59 cell supernatant was diluted to the desired concentration and mixed with human whole blood provided by the American Red Cross (IBC protocol #2010-014), taking care to ensure that the virus solution added to the blood did not exceed 10% of the total volume to minimize dilution of the blood’s nutritional value. The final titer of the blood meal for both trials was 6.40e07 viral copies/mL as verified by qPCR and 1.3e03 PFU/mL as determined by plaque assay, which corresponds with the clinically observed range of ZIKV in human blood titers. *(28)* Thirty 2-day old adult female mosquitoes were transferred to small containers and allowed to feed on the ZIKV supplemented blood from a membrane feeding system. After feeding for 1 hour, mosquitoes were cold anaesthetized and female mosquitoes with visibly engorged abdomens were separated, while non-blood-fed mosquitoes were discarded. A total of 180 blood-fed female mosquitoes, 20 mosquitoes/cage, were separated into three temperature treatments of 27°C, 30°C, and 33°C with 60 female mosquitoes (i.e. 3 cages) per treatment. At 0 days post infection, each cage was provided with an oviposition substrate and female mosquitoes were allowed to oviposit freely for 72 hours, representing the first gonotrophic cycle, after which the egg sheet was removed. Maternal mosquitoes were provided additional uninfected blood meals at 5 and 10 days post infection (dpi) to facilitate additional reproductive cycles. Oviposition substrates were provided after each blood meal, corresponding to the second and third gonotrophic cycles.

### Vertical Transmission to Progeny

Potentially vertically-infected progeny of orally-infected female mosquitoes were reared according to the methods described above. After adult emergence, progeny mosquitoes were maintained in small cages at 20 mosquitoes/cage. At 3dpi, 7 dpi, and 14 dpi, ten adult female mosquitoes from each maternal temperature treatment and gonotrophic cycle cohort were cold anesthetized to collect saliva and abdomen samples. Legs and wings were removed, then each female’s proboscis was inserted into a 0.2 mm capillary tube and allowed to salivate into mineral oil for five minutes, after which the saliva sample was aspirated into 200mL of DMEM mixed with 2% FBS, 1% pen/strep for processing. Saliva was tested for viral presence as a measure of potential infectiousness to humans and stored in −80°C for RNA extraction. Abdomen samples were collected in the same way as the I1s.

Leg/wing and abdomen samples were homogenized in 1.6 mL tubes with 500 uL of Trizol and Total RNA isolated with Trizol as described above. For saliva samples, 100 ul of saliva/DMEM solution was placed in 200uL of Trizol to inactivate any virus, then processed as above. Concentrations of the extracted RNA were verified using a Nanodrop 2000 (ThermoFisher, Waltham MA) and stored in −80°C until a viral titer could be determined by qPCR as described above. To verify successful RNA extraction and cDNA synthesis, a qPCR reaction was run with *Ae. aegypti* actin primers and SYBR Green one step kit according to manufacturer protocol (ThermoFisher, Waltham MA).

### Statistical Analyses

Evaluating the effects of the explanatory variables (i.e., maternal post-infection temperature, gonotrophic cycle, and progeny age) simultaneously using logistic regression was not possible because the sample size did not allow this type of stratification of the data and resulted in unstable estimates of the regression coefficients. Accordingly, separate univariate logistic regression models were used to evaluate the effect of 1) maternal post-infection temperature, 2) gonotrophic cycle, 3) progeny age, and 4) experimental trial (n=2) on the odds of progeny infection. Experimental trial was treated as a categorical variable, while maternal temperature (27°C, 30°C, or 33°C), gonotrophic cycle (first, second, or third), and progeny age (3, 7, or 14 days post-eclosion) were treated as numerical variables. Three outcomes were modeled: odds of progeny with detectable virus overall; odds of progeny with detectable virus in saliva; and odds of progeny with detectable virus in the abdomen. Exact binomial 95% confidence intervals were calculated for the percentage of ZIKV-infected mosquitoes for each level of the explanatory variables. Odds ratios reported for numerical predictors are unit odds ratios. For the categorical predictor experimental trial, the first trial was held as the reference category and the odds of the second trial relative to the first are reported. A Pearson chi-squared test with Yates continuity correction was used to evaluate whether viral presence in the abdomen was associated with virus presence in the saliva of individual mosquitoes. The observed proportion of progeny mosquitoes with both infected abdomens and saliva was compared to the expected proportion if the two types of infection are independent, which was the product of the observed proportions of progeny mosquitoes with infected abdomens only and infected saliva only.

## Results

### *Orally-infected* Ae. aegypti *females transmit ZIKV to their offspring*

To assess the rate of vertical transmission, we examined the offspring of orally-infected female *Ae. aegypti* for ZIKV presence in abdomens and saliva at multiple time points over a two week period post-eclosion. Vertical transmission from orally-infected *Ae. aegypti* females to their offspring occurred across all maternal temperature conditions, gonotrophic cycles, and adult progeny ages tested. Overall, 6.5% (95% confidence interval = 3.9 - 9.9) of adult progeny had detectable levels of ZIKV. Experimental trial was not a significant explanatory variable in any analysis, indicting independence of progeny outcome from experimental trial. (Table 1)

**Table 1:** Logistic Regression Analysis of ZIKV Vertical Transmission. Each unique combination of explanatory variable and progeny infection outcome represents a distinct univariate regression model. Odds ratios for numerical explanatory variables are reported as unit odds ratios, odds ratios for categorical predictors are relative to the reference category. Statistically significant models (p ≤ 0.05) are in bold.

| Explanatory Variable | Progeny Infection Outcome | X<sup>2</sup> | p-value | Odds Ratio |
| --- | --- | --- | --- | --- |
| *Experimental Trial* | Total | 0.92 | 0.34 | 0.59 |
|  | Abdomen | 1.41 | 0.24 | 0.39 |
|  | Saliva | 0.11 | 0.74 | 0.79 |
| *Maternal Temperature* | Total | 1.59 | 0.21 | 0.87 |
|  | Abdomen | 0 | 0.99 | 1 |
|  | Saliva | 2.72 | 0.10 | 0.75 |
| *Gonotrophic Cycle* | Total | **3.84** | **0.05** | **1.95** |
|  | Abdomen | 1.76 | 0.18 | 1.92 |
|  | Saliva | 1.19 | 0.28 | 1.76 |
| *Progeny Age* | Total | **3.91** | **0.05** | **0.87** |
|  | Abdomen | **4.07** | **0.04** | **0.77** |
|  | Saliva | 0 | 0.99 | 1 |

### ZIKV vertical transmission occurs at all maternal temperature conditions

To assess the effect of maternal temperature on vertical transmission we maintained orally-infected maternal mosquitoes at 27°C, 30°C, and 33°C. ZIKV was present in adult progeny from every maternal temperature. Vertical transmission occurred to 8.3% (n = 109, 95% CI = 3.8 - 15.1) of progeny from 27°C, 5.7% (n = 105, 95% CI = 2.1 - 12) from 30°C, and 3.8 % (n = 79, 95% CI = 0.8 - 10.7) from 33°C. (Figure 1A) ZIKV was found in progeny abdomens at every temperature; 4% (n = 100, 95% CI = 1-1 – 9.9) of progeny abdomens from 27°C, 3% (n = 99, 95% CI =0.6 - 8.6) from 30°C, and 3.8% (n = 79, 95% CI = 0.8 – 10.7) from 33°C. ZIKV was found in the saliva of 4.9% (n = 102, 95% CI = 1.6 – 11.1) of progeny from 27°C and 3.8% (n = 104, 95% CI = 1.5 – 9.6) from 30°C, but not in any of the 77 progeny from the 33 °C treatment. (Figure 1B) There was no significant association between maternal temperature and the odds of ZIKV infection in the progeny mosquitos. (Table 1)

**Fig 1:**
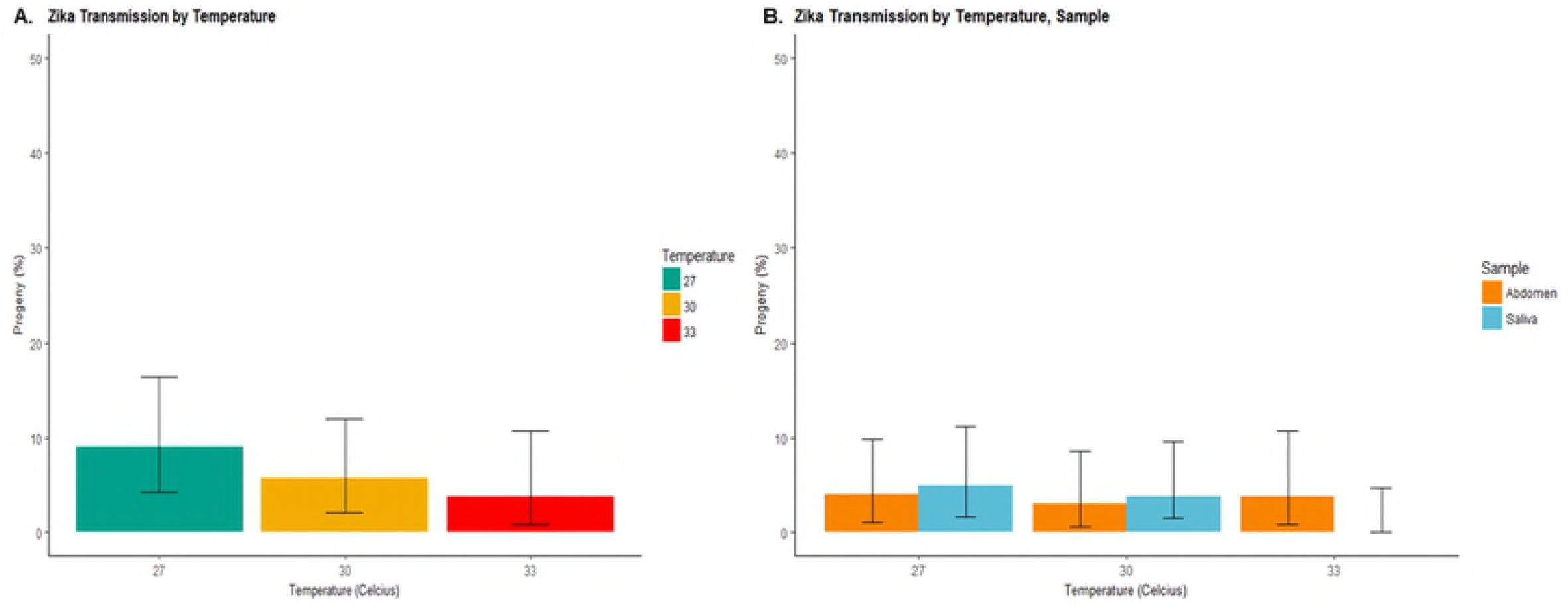
Vertical Transmission by Maternal Temperature and Sample Type. (A) Percent of vertically infected progeny from each maternal temperature condition. (B) Percent of progeny with ZIKV present in abdomens, indicating vertical infection, and saliva, indicating potential infectiousness, from each maternal temperature condition.

### ZIKV Vertical transmission occurs at every gonotrophic cycle

To determine whether vertical transmission increased as the time between maternal infection and oviposition increased we assessed the rate of vertical transmission over consecutive gonotrophic cycles. Vertical transmission occurred in every gonotrophic cycle. ZIKV was detected in 4.0% (n = 174, 95% CI = 1.6 - 8.1) of progeny from the first gonotrophic cycle, 8.6% (n = 105, 95% CI = 4 - 15.6) from the second, and 14.3% (n = 14, 95% CI 1.8 - 42.8) from the third. (Figure 2A) ZIKV was present in the abdomens of 1.7% (n = 174, 95% CI = 0.4 – 5.0) progeny from the first gonotrophic cycle and 7.8% (n = 90, 95% CI = 3.2 – 15.4) progeny from the second but was not detected in the abdomens of the fourteen progeny from the third gonotrophic cycle. ZIKV was detected in the saliva of 3.0% (n = 166, 95% CI = 1.0 – 6.9) progeny from the first cycle, 1.9% (n = 104, 95% CI = 0.2 – 6.8) of progeny from the second cycle, and 16.7% (n = 12, 95% CI = 2.1- 48.4) of progeny from the third cycle. (Figure 2B) The relatively small sample size of the third gonotrophic cycle was due to lower survivorship and fecundity of maternal mosquitoes after laying two previous batches of eggs. Gonotrophic cycle significantly affected the odds of overall ZIKV infection in progeny (p = 0.05, Odds ratio = 2), suggesting a greater number of developing ovarioles became infected with each subsequent cycle. However, there was no discernable influence of gonotrophic cycle on the odds of viral presence in abdomen tissue or saliva samples individually. (Table 1)

**Fig 2:**
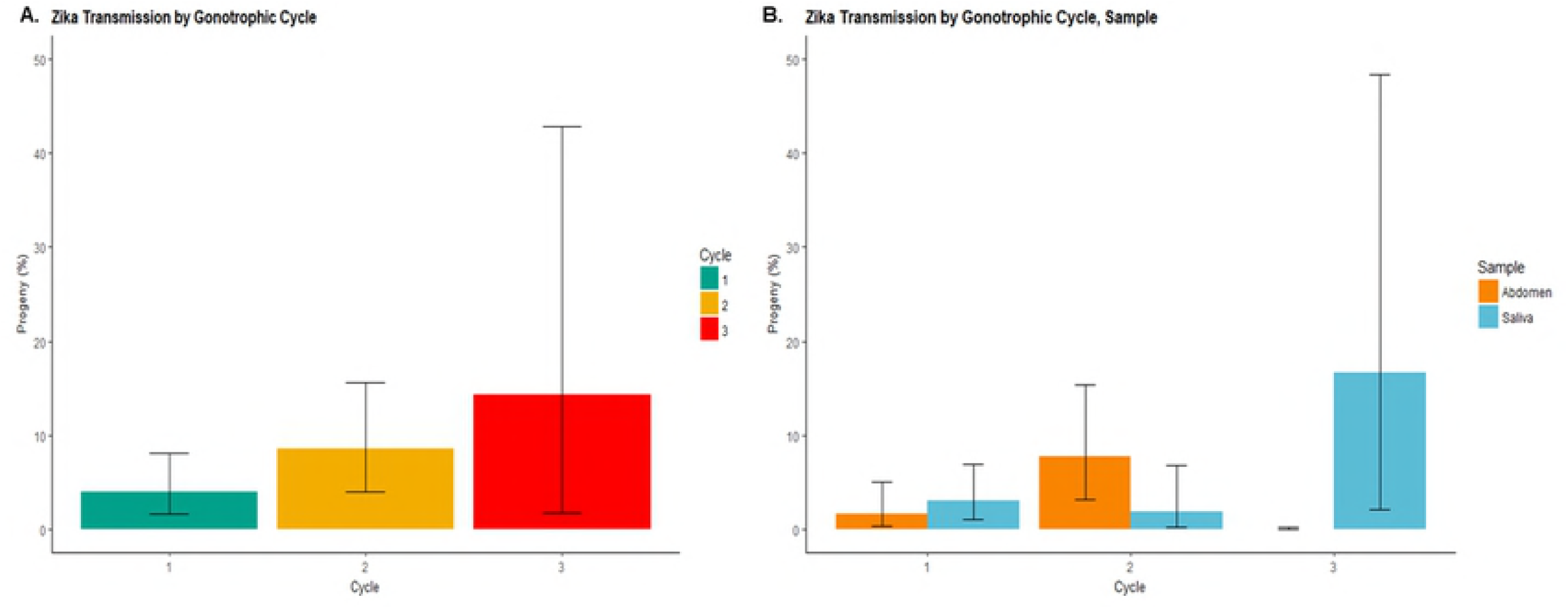
Vertical Transmission by Gonotrophic Cycle and Sample Type. (A) Percent of vertically infected progeny from each gonotrophic cycle. (B) Percent of progeny with ZIKV present in abdomens, indicating vertical infection, and saliva, indicating potential infectiousness, from each gontrophic cycle.

### ZIKV infection persists for at least two weeks in infected progeny

Persistence of ZIKV in vertically infected progeny is essential for viral maintenance within mosquito populations and would result in greater opportunities for the virus to be passed to vertebrate hosts during bloodfeeding. We found ZIKV in progeny up to two weeks after adult eclosion. ZIKV was detected in 11.7 % (n = 103, 95% CI = 6.2 - 19.5) of three day old progeny, 2.8 % (n = 108, 95% CI = 0.6 - 7.9) of seven day old progeny, and 3.7 % (n = 82, 95% CI = 0.8 - 10.3) of fourteen day old progeny. (Figure 3A) ZIKV was detected in the abdomens of 8.2% (n = 98, 95% CI = 3.6 – 15.5) of three day old progeny, 1.0% (n = 98, 95% CI = 0 – 5.6) of seven day old progeny, and 1.2% (n = 82, 95% CI = 0 – 6.6) of fourteen day old progeny. ZIKV was detected in the saliva of 4.4% (n = 91, 95% CI = 1.2 – 10.8) of three day old progeny, 1.9% (n = 105, 95% CI = 0.2 – 6.7) of seven day old progeny, and 3.4% (n = 87, 95% CI = 0.8 – 9.7) of fourteen day old progeny. (Figure 3B) Progeny age was significantly associated with reduced odds of virus presence in progeny mosquitoes overall (p = 0.05, Odds ratio = 0.87) and in progeny abdomens (p = 0.04, Odds ratio = 0.77), but not with viral presence in saliva. (Table 1)

**Fig 3:**
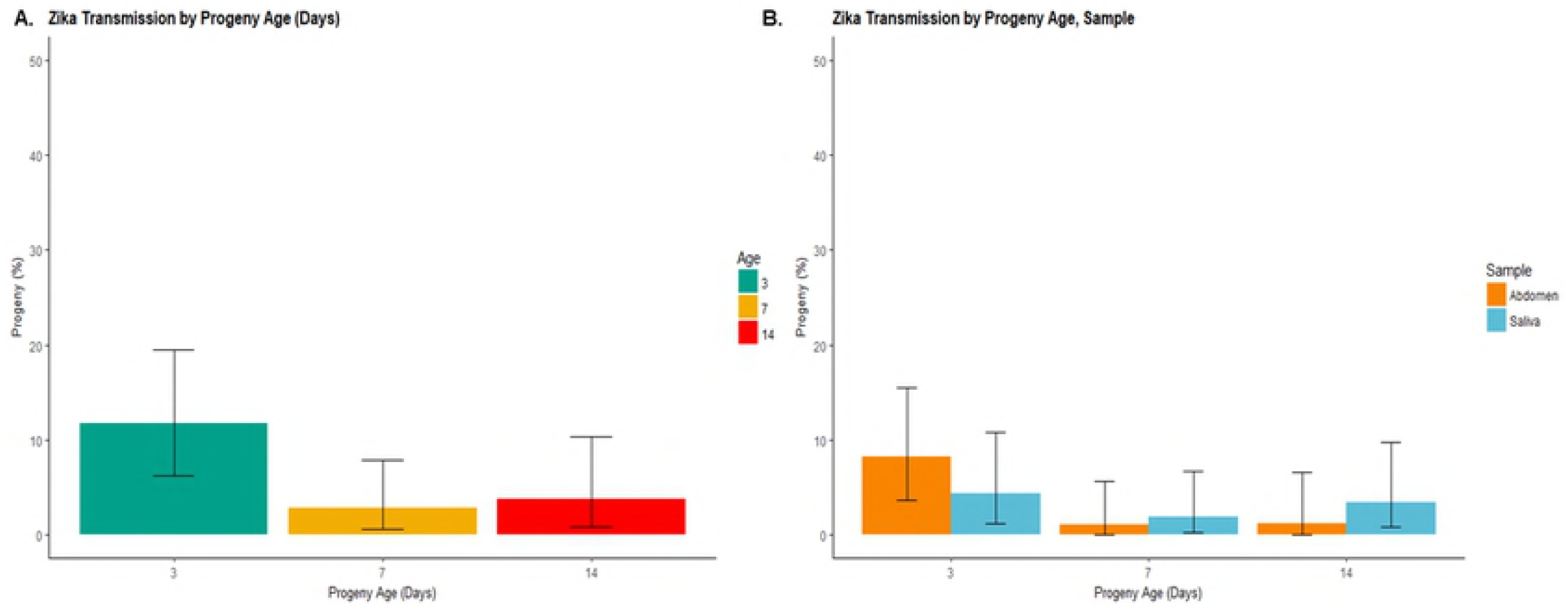
Vertical Transmission by Progeny Age and Sample Type. (A) Percent of vertically infected progeny by age (days) at dissection. (B) Percent of progeny with ZIKV present in abdomens, indicating vertical infection, and saliva, indicating potential infectiousness, by age (days) at dissection.

### Vertically infected progeny have ZIKV in their saliva

For transmission to the vertebrate host to occur, virus must be present in the mosquito’s saliva. ZIKV positive saliva was detected in 3.4 % (1.6 - 6.2) of progeny, or half of all vertically-infected progeny. Virus was found in saliva of adult progeny hatched from every gonotrophic cycle and at every progeny age tested. ZIKV positive saliva was detected in progeny from all maternal temperature conditions except 33°C, though the sample size was smaller for this temperature (n = 77). None of the explanatory variables were significantly associated with ZIKV presence in saliva. (Table 1) Additionally, though virus was detected in just the saliva or just the abdomens of 3.07% of progeny mosquitoes, viral presence in progeny abdomens was not significantly associated with viral presence in saliva (X^2^ = 0.001, p = 0.97), and only occurred in one progeny mosquito.

## Discussion

Understanding the capacity of ZIKV to establish itself in vector mosquito populations, persist through periods of low transmission, and initiate future outbreaks is vitally important to public health. We found vertical transmission occurred from orally infected female *Ae. aegypti* to 6.5% (95% CI = 3.9 - 9.9) of adult female progeny. This is consistent with vertical transmission rates of other Aedes-borne flaviviruses, including DENV-3 (3%), chikungunya (20.2%), and yellow fever (8.2%) (*29, 11, 12*) and is substantiated by evidence from the field. ZIKV positive males have been identified in Mexico and Brazil, confirming its occurrence. (4,33) Whether this rate of vertical transmission is sufficient to contribute to future outbreaks has been debated. The estimated minimum rate of vertical transmission required to influence human cases of a similar arbovirus, dengue, ranges from 4% to 20-30%. (*30, 31*) Some studies have contested that human movement and asymptomatic cases contribute more significantly to human dengue prevalence than vertical transmission. (31, 32) However, peaks in the vertical transmission of dengue have been shown to precede disease outbreaks during periods of high mosquito density. *(34)* At a minimum, vertical transmission should be considered as one of ZIKV’s multiple modes of transmission and incorporated into prevention strategies, surveillance plans, and models of transmission risk.

Vertical transmission of ZIKV throughout an infected mosquito’s reproductive lifetime could facilitate pathogen establishment. Considering all progeny tested, orally-infected *Ae. aegypti* females were twice as likely to vertically transmit ZIKV to their progeny with each consecutive gonotrophic cycle (p = 0.05, Odds ratio = 2), suggesting a greater number of ovarioles were invaded by virus as the maternal mosquito aged. However, analyses considering abdomen and salivary infection separately did not find a significant association between gonotrophic cycle and progeny infection (Table 1), which could have arisen because of reduced statistical power due to the low number of progeny infected overall. Mosquito survivorship in the field is significantly lower than under laboratory conditions, and even in the lab not many maternal mosquitoes lived to complete a third gonotrophic cycle, as reflected by our small sample size. *(16)* Accordingly, the shorter reproductive lifetime of mosquitoes in the field may attenuate the impact of higher transmission rates during later gonotrophic cycles.

Vertical transmission occurred across a wide range of maternal temperature conditions, suggesting that it could be a robust mechanism for ZIKV maintenance in mosquitoes. During the 2015 pandemic, ZIKV spread to a variety of climates and this window of temperatures suitable for vertical transmission could be one of opportunity for ZIKV establishment. (1) Geographic regions with a competent vector population and average temperatures that overlap 27-33°C, including much of the southern United States, are at risk of ZIKV vertical transmission. (35) Interestingly, ZIKV vertical transmission was not significantly associated with higher temperatures, unlike horizontal transmission of a similar arbovirus; dengue. (*22*)

Persistence of ZIKV vertical infection up to two weeks after adult progeny eclosion extends the possibility of pathogen persistence in previous outbreak areas and transmission to human hosts. However, lifelong presence of ZIKV in the saliva of progeny mosquito, as opposed to saliva infection of older mosquitoes with horizontal transmission, may alter aspects of the virus-vector relationship. In horizontally infected mosquitoes, virus must be acquired from an infected host, which may not occur during the first bloodfeeding. The virus must then disseminate through the body, including the abdomen, to reach the salivary glands. (15) However, for vertical transmission, ZIKV presence in the abdomens of infected progeny appeared independent of presence of ZIKV in the saliva (p = 0.97). The relationship between mosquito age and likelihood of viral infection also differs between horizontal and vertical transmission. Vertically infected progeny were less likely to test positive for ZIKV as they aged, with a 13% reduction in the odds of overall infection (p = 0.05, Odds ratio = 0.87) and a 23% reduction in the odds of abdomen infection (p = 0.04, Odds ratio = 0.77). This relationship is reversed in horizontally infected mosquitoes, as dengue transmission increases with mosquito lifespan. (*22*)

The presence of virus in the saliva of vertically infected progeny provides ZIKV with a potential bridge between mosquitoes and humans. This bridge was present at all progeny ages tested, with ZIKV detected in the saliva as early as three days and as late as 14 days after adult emergence. Whether there is a difference in vector competence between vertically and horizontally infected mosquitoes should be the subject of further research. Not only do vertically-infected progeny have the capacity to start an outbreak, they likely have greater opportunity to do so than horizontally-infected mosquitoes. Vertically infected mosquitoes bypass two essential requirements necessary for horizontal transmission, obtaining the initial infectious blood meal and surviving the extrinsic incubation period (EIP). The maternal mosquitos in our study had a minimum EIP of 3 days *(R. Zinna, unpub. data)*, and other studies found the average ZIKV EIP to range from 4 to 10 days. (*14,15*) Without the need to find an initial infectious bloodmeal and survive the multi-day waiting period imposed by the EIP, vertically infected adult progeny are capable of transmitting the virus to humans at a younger age and for a greater percentage of their adult lives.

### Conclusion

We found vertical transmission from orally infected females occurs as early as the first reproductive cycle following the initial infectious bloodmeal, at a range of maternal temperatures, and that ZIKV infection persists in the progeny for at least two weeks after adult eclosion. ZIKV was detected in the saliva of vertically infected progeny, and the resulting elimination of the extrinsic incubation period means these mosquitoes have a longer interval than horizontally infected mosquitos to bite and infect a human. Consequently, ZIKV has the potential to be maintained in mosquito populations even in the absence of transmission cycles to the vertebrate host and thus can cause future outbreaks without the need for viral reintroduction. These findings reinforce the need to conduct surveillance and viral testing in mosquito populations where ZIKV transmission has occurred in the past.

## Acknowledgements

The authors would like to extend a heartfelt thank you to Teresa Joy, for her assistance in the lab, Michael Pham, for his cell-culture expertise, and Jenet Soto-Shoumaker, for maintaining the large research colonies that made this work possible. We would also like to thank Kathryn Fitzpatrick for her troubleshooting advice.

